# Auto-inhibitory regulation of DNA binding by the C-terminal tails of the mitochondrial transcription factors Mtf1 and TFB2M

**DOI:** 10.1101/2020.03.06.980961

**Authors:** Urmimala Basu, Nandini Mishra, Mohammed Farooqui, Jiayu Shen, Laura C. Johnson, Smita S. Patel

**Author notes:** To whom correspondence should be addressed: Smita S. Patel: Department of Biochemistry and Molecular Biology, Rutgers University, Robert Wood Johnson Medical School, Piscataway, NJ 08854, USA.

## Abstract

The structurally homologous Mtf1 and TFB2M proteins serve as transcription initiation factors of the *Saccharomyces cerevisiae* and human mitochondrial RNA polymerases, respectively. These transcription factors directly interact with the non-template strand of the transcription bubble to drive promoter melting. Given the key roles of Mtf1 and TFB2M in promoter-specific transcription initiation, it is expected that the DNA binding activity of the mitochondrial transcription factors would be regulated to prevent DNA binding at inappropriate times. However, there is little information on how mitochondrial DNA transcription is regulated. While studying the C-tail deletion mutants of Mtf1 and TFB2M, we stumbled upon a new finding that suggested that the flexible C-tail region of these factors autoregulates their DNA binding activity. Quantitative DNA binding studies with fluorescence anisotropy-based titrations show that Mtf1 with an intact C-tail has no affinity for the DNA but the deletion of C-tail greatly increases the DNA binding affinity. Similar observations were made with TFB2M, although autoinhibition by the C-tail of TFB2M was not as absolute as in Mtf1. Analysis of available TFB2M structures show that the C-tail makes intramolecular interactions with the DNA binding groove in the free factor, which we propose masks the DNA binding activity. Further studies show that the RNA polymerase relieves autoinhibition by interacting with the C-tail and engaging it in complex formation. Thus, our biochemical and structural analysis identify previously unknown autoinhibitory and activation mechanisms of mitochondrial transcription factors that regulate the DNA binding activity and aid in specific assembly of the initiation complexes.

## Introduction

Mitochondrial RNA polymerases (RNAPs) are evolutionarily related to single-subunit bacteriophage T7/T3 RNAP (1). However, unlike T7/T3 RNAP, the core subunit of the mitochondrial RNAP is unable to initiate promoter-specific transcription and requires assistance from accessory transcription initiation factors. To catalyze promoter-specific transcription initiation, the core subunit of *Saccharomyces cerevisiae* (yeast) mitochondrial RNAP, Rpo41, requires Mtf1 (Mitochondrial Transcription Factor 1) (2,3) and the human mitochondrial RNAP, POLRMT, requires TFB2M (Transcription Factor B2 Mitochondrial) and TFAM (Transcription Factor A Mitochondrial) (3-7) These factors play a key role in stabilizing the transcription initiation bubble.

Mtf1 and TFB2M proteins are evolutionarily related to bacterial rRNA methyltransferases (8,9). The methylation activity is not required for transcription initiation; hence, Mtf1 has lost the methyltransferase activity, but interestingly, TFB2M has residual methyltransferase activity (10). However, both these factors have maintained their nucleic acid binding function, which is essential for promoter melting. The nucleic acid binding groove in the factors serves as a binding pocket for the non-template strand to stabilize the transcription initiation bubble (8,11).

Transcription factors are regulated in various ways, including reversible protein phosphorylation, accessory factors, and by autoregulatory mechanisms (12-16). For example, bacterial sigma factors are regulated by anti-sigma factors or autoinhibited by the sigma-1.1 domain (16). No such regulatory mechanism has been reported for the mitochondrial transcription factors.

During our studies of the C-terminal deletion mutants of Mtf1 (3), we stumbled upon a new finding that suggested that the flexible C-terminal tail of Mtf1 has an autoinhibitory function in DNA binding. Both Mtf1 and TFB2M contain a flexible C-tail region consisting of 16 to 20 aa, which we recently showed plays a critical role in template strand alignment including supporting high affinity binding of the initiating nucleotide for efficient RNA priming reaction (3). The 16 aa of the Mtf1 C-tail were not resolved in the crystal structure (9), whereas in the crystal structures of TFB2M, the fully or partially resolved C-tail region was found to be in different conformations (8).

To investigate the regulatory role of the flexible C-tail region in DNA binding, we used fluorescence anisotropy-based titrations and measured the DNA *K*_d_ values of WT and several C-tail deletion mutants of Mtf1 and TFB2M. We used both promoter and non-promoter sequences to test the specificity of the DNA binding site. Our studies show that DNA binding is non-specific and the C-tail region completely or partially suppresses the DNA binding activity of the factors. Inhibition is released, however, when the factors bind to the RNAP and the C-tail gets engaged with the RNAP. Based on our biochemical data and analysis of available structures, we propose a model that explains the autoinhibition and activation of DNA binding involving the C-tail of the mitochondrial transcription factors. This model provides new insights into the assembly and regulation of the initiation complex.

## Results

### The C-tail inhibits the DNA binding activity of Mtf1

In our studies, we used two Mtf1 C-tail deletion mutants which were used in our previous studies (3). Mtf1-Δ20 lacks the entire C-tail region (20 aa) and the partially deleted C-tail mutant Mtf1-Δ12 lacks only the terminal 12 aa (Fig. 1A). Because Mtf1 binds to the non-template strand of the transcription bubble in the initiation complex (11), we synthesized a 12-nt single-stranded DNA that contained the −8 to +4 sequence of the non-template strand of the yeast *15S* promoter (Fig. 1B). This DNA included the - 4 to +2 region that forms a transcription bubble (17) and interacts with Mtf1 in the initiation complex (11).

**Fig. 1:**
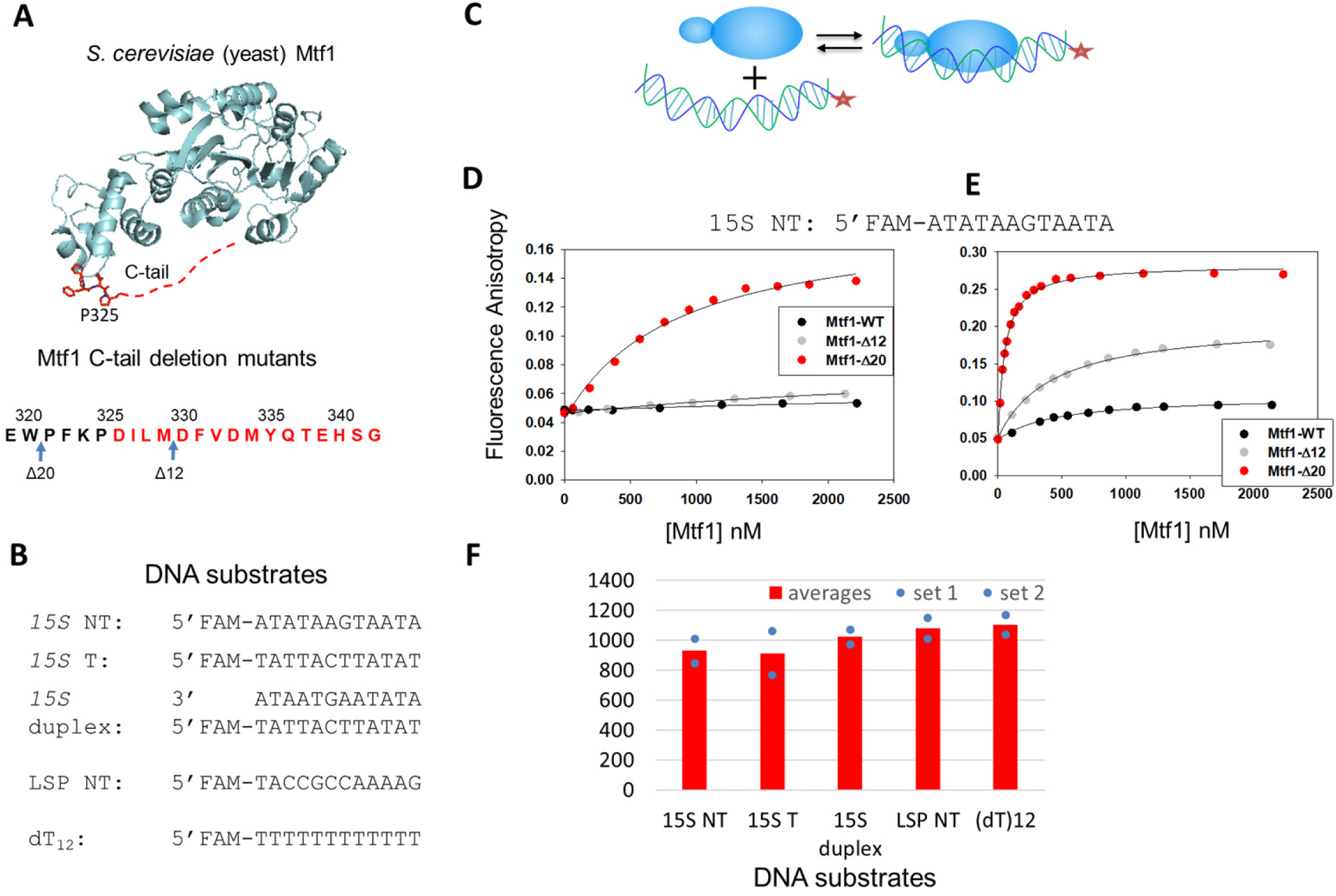
The C-tail of Mtf1 drastically autoinhibits the DNA binding activity of Mtf1. (A) Structure of the yeast mitochondrial transcription factor Mtf1 in grey (PDB ID:1i4w). The missing 16 aa of the C-tail of Mtf1 in the crystal structure are shown in red dotted line and also marked in red in the amino acid sequence of the C-tail of Mtf1. (B) DNA sequences of the substrates used for the Mtf1 DNA binding studies. (C) Cartoon showing the basic scheme of the fluorescence anisotropy assays to monitor protein-DNA binding. (D) Representative binding curves showing the fluorescence anisotropy changes resulting from titration of the *15S* NT DNA with Mtf1. *15S* NT DNA (5 nM) was titrated with Mtf1-WT (black circles), Mtf1-Δ12 (grey circles) and Mtf1-Δ20 (red circles) in Buffer A (Experimental procedures). (E) *15S* NT (5 nM) was titrated with Mtf1-WT (black circles), Mtf1-Δ12 (grey circles) and Mtf1-Δ20 (red circles) in Buffer A without potassium glutamate. The solid lines represent fit to the hyperbolic equation with *K*_d_ values as follows: Mtf1-WT = 447±60 nM (amplitude: 0.059); Mtf1-Δ12 = 426±33 nM (amplitude: 0.156); Mtf1-Δ20 = 51±2.8 nM (amplitude: 0.24). The errors represent standard error of the fit. (F) The average DNA *K*_d_ values of Mtf1-Δ20 are shown for the DNA substrates in (B). The blue dots are the individual values for set 1 and set 2 titration data which are shown in Figure S1.

We used fluorescence anisotropy-based titrations to measure the DNA *K*_d_ values (Fig. 1C) (18). Fluorescein labeled *15S* NT was titrated with increasing concentrations of Mtf1 protein. We observed negligible changes in fluorescence anisotropy with Mtf1-WT and Mtf1-Δ12 even after adding 2 µM protein (Fig. 1D). On the other hand, Mtf1-Δ20, with a larger deletion of the C-tail, showed a significant increase in fluorescence anisotropy in the titration experiments. The binding data of Mtf1-Δ20 fit well to a hyperbola and provided a *K*_d_ of ∼870 nM for the *15S* NT complex. These results indicate that the presence of the full-length or even partial C-tail region inhibits the DNA binding activity of Mtf1. Thus, a complete deletion of the C-tail is needed to activate the DNA binding activity of Mtf1.

The DNA binding buffer in the above experiments contained 50 mM potassium glutamate. To determine if the DNA binding interactions were salt sensitive, we eliminated potassium glutamate from the buffer. Under less stringent salt conditions, we observed some amount of DNA binding to both the WT protein and Mtf1-Δ12, although the amplitudes of fluorescence anisotropy change remained lower than with Mtf1-Δ20 (Fig. 1E). Interestingly, there was a clear C-tail length dependent effect on the DNA binding amplitudes. Mtf1-WT with an intact C-tail showed the lowest amount of DNA binding, followed by Mtf1-Δ12 with an intermediate level of binding, and Mtf1-Δ20 showing the highest amount of DNA binding. Removal of the salt increased the DNA binding affinity of Mtf1-Δ20 by 18-fold. These results point to an ionic nature of interaction in the DNA binding groove and suggest that Mtf1 makes contact with the charged DNA phosphate backbone. Thus, competition with salt explains the lower DNA binding affinity of Mtf1-Δ12.

To investigate whether the DNA binding groove of Mtf1 has a preference for binding a particular DNA sequence or structure, we compared the DNA *K*_d_ values of promoter versus non-promoter sequences and single-stranded versus double-stranded DNAs. We used both the template and non-template strands of the yeast *15S* promoter as DNA substrates, the duplex *15S* promoter, an unrelated DNA sequence of the human mitochondrial promoter (LSP NT), and dT_12_ (Fig. 1B). Mtf1-WT did not bind to any of these DNA substrates. Mtf1-Δ20, on the other hand, bound to all the DNA substrates (Fig. 1F & S1) with very similar *K*_d_ values (750-1000 nM). These results indicate that the DNA binding groove of Mtf1 has no sequence or structural preference for the DNA. This means that the C-tail prevents Mtf1 from binding to both specific and non-specific DNAs.

In summary, the above described DNA binding studies provide the first evidence that the C-tail region has a role in regulating the DNA binding activity of Mtf1 prior to transcription initiation.

### Structural basis for the C-tail mediated inhibition of DNA binding

To understand how the C-tail region of Mtf1 inhibits DNA binding, we analyzed the structures of the homologous human TFB2M protein (8). The C-tail region in free TFB2M is fully resolved in chain A of the molecule (Fig. 2A, TFB2M in green and C-tail in red). In chain A, the C-tail is making intramolecular interactions with the DNA binding groove (Fig. S2). In the initiation structure, the DNA binding groove is engaged with the non-template strand (in blue); and therefore, the C-tail (in orange) is projected in a different direction away from the DNA binding groove (Fig. 2A). Thus, analysis of the TFB2M structures suggests that the conformation of the flexible C-tail in free TFB2M is sterically incompatible with DNA binding. Although Mtf1 and TFB2M share only 12% sequence identity, there is a high structural homology between the two proteins (Fig. 2A, Mtf1 in pink and TFB2M in green). We propose that the DNA binding activity of Mtf1 is similarly autoinhibited by C-tail interactions with the DNA binding groove.

**Fig. 2:**
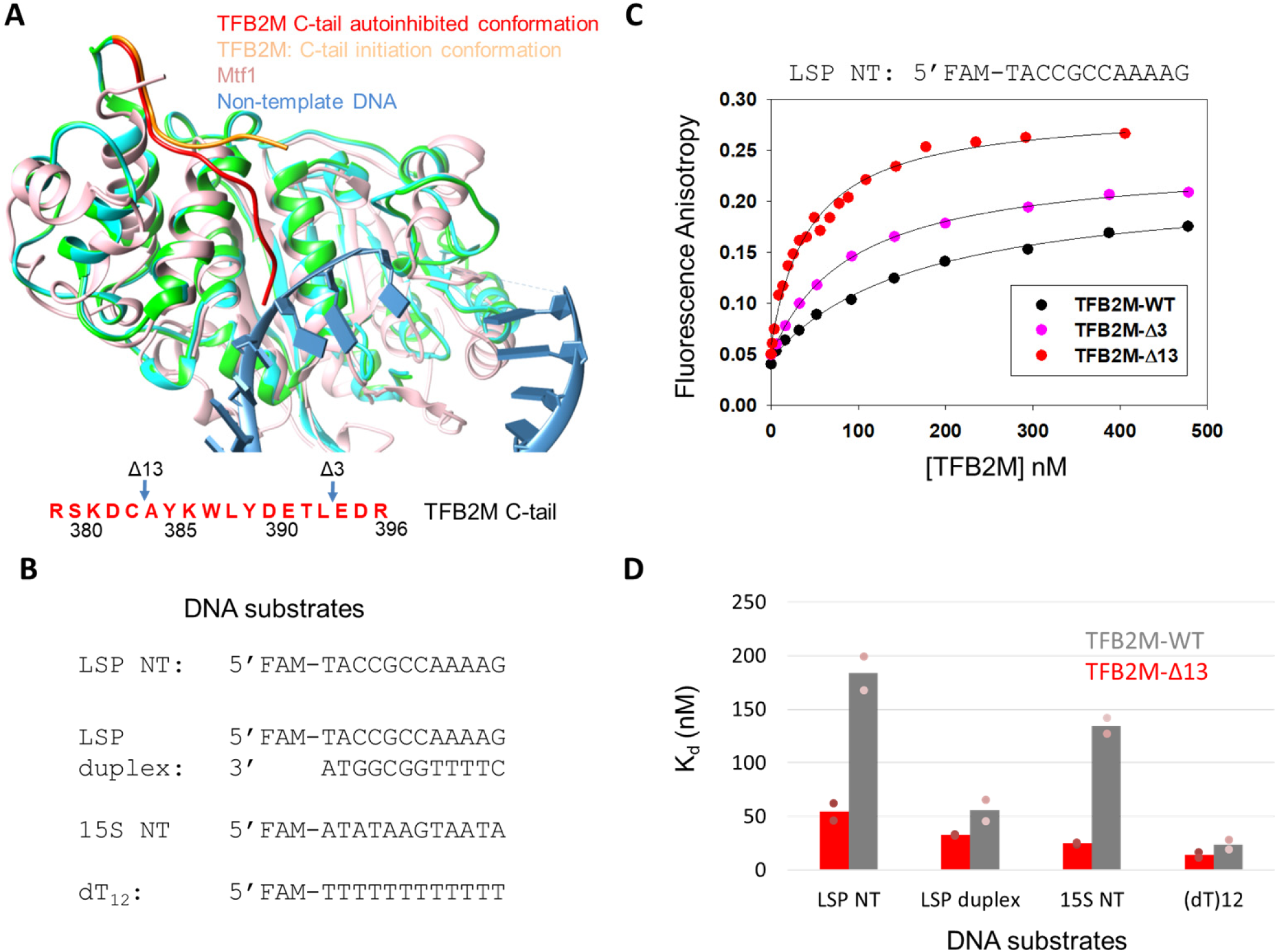
The C-tail of TFB2M mildly autoinhibits the DNA binding activity of TFB2M. (A) The aligned structures of free human TFB2M (PDB: 6ero in cyan), TFB2M in the initiation complex (PDB: 6erp in green), and the yeast Mtf1 (PDB: 1I4W in white) are shown. The relative positions of the C-tail in all three structures are shown. The amino acid sequence of the C-tail of TFB2M is shown below in red. (B) DNA substrates used for the TFB2M DNA binding studies. (C) Representative binding curves show the fluorescence anisotropy change resulting from titration of LSP NT (5 nM) with TFB2M-WT (black circles), TFB2M-Δ3 (pink circles), and TFB2M-Δ13 (red circles). The data were fit to a hyperbola to obtain the following *K*_d_ values: TFB2M-WT = 169±18 nM (amplitude 0.17); TFB2M-Δ3 = 92±3.3 nM (amplitude 0.19); TFB2M-Δ13: 46±5.2 nM (amplitude 0.23). (D) The red and grey bars show the DNA *K*_d_ values of TFB2M-WT and TFB2M-Δ13, respectively, for the various DNA substrates shown in (B). The blue dots represent individual values for set 1 and set 2 titration data that are shown in Figure S3.

### Autoinhibitory role of the C-tail is conserved in the human homolog TFB2M

The above structural analysis predicts that the C-tail will inhibit the DNA binding activity of TFB2M. To directly test the autoinhibitory role of the C-tail in TFB2M, we measured the DNA *K*_d_ values of TFB2M-WT and its two C-tail deletion mutants, TFB2M-Δ3 and TFB2M-Δ13 that lacks 3 and 13 aa of the C-tail, respectively (Fig. 2A) (3). In our initial DNA binding studies, we used the non-template strand of the human mitochondrial light strand promoter (LSP NT) as the substrate, which contains the DNA sequence that binds to TFB2M in the initiation complex. Additionally, we synthesized the LSP duplex, and unrelated *15S* NT and dT_12_ to determine whether TFB2M has a preference for binding to promoter sequences (Fig. 2B).

TFB2M-WT binds to LSP NT with a *K*_d_ of about 180 nM and deletion of 13 aa of the TFB2M C-tail increases the DNA binding affinity of LSP NT by about 4-fold (Fig. 2C & S3). Deletion of 3 aa of the C-tail results in intermediate level of DNA binding (Fig. 2C, Fig. S3). This shows a C-tail length dependent effect on the DNA binding activity, similar to Mtf1.

Studies with other DNA sequences indicate that TFB2M binds to all the DNAs but with different *K*_d_ values (Fig. 2D). The non-promoter *15S* NT binds with a *K*_d_ of 130 nM, which is similar to the *K*_d_ of LSP NT. Thus, DNA binding activity of free TFB2M appears to be non-specific. The LSP duplex has a 3-fold higher affinity than the LSP NT. For reasons unknown, TFB2M has a high affinity for dT_12_ DNA. In all cases, however, C-tail deletion increases the DNA binding affinity of TFB2M.

Thus, our results show that the C-tail of TFB2M autoinhibits the DNA binding activity but autoinhibition by the C-tail of TFB2M is not absolute as observed in Mtf1. The general trend is the same; that is, the presence of the C-tail inhibits the DNA binding activity. Thus, the C-tail has a conserved role in autoinhibiting the DNA binding activity of the free mitochondrial transcription factors.

### The C-tail of Mtf1 is required for stable complex formation with the RNAP subunit

Next, we asked how the DNA binding activity of Mtf1 and TFB2M was activated for transcription initiation. Structural analysis shows that in the initiation complex the partially resolved C-tail of TFB2M interacts with the RNAP subunit, POLRMT (Fig. S4) (8). This suggests that RNAP may release the autoinhibited state by engaging the C-tail in an altered conformation.

To investigate the RNAP mediated mechanism of activation, we investigated whether the C-tail has a role in complex formation with the RNAP subunit. We used Bio-layer Interferometry (BLI) and ultrafiltration approaches and measured protein-protein interactions between Mtf1 and the RNAP subunit, Rpo41. His-tagged Mtf1-WT and Mtf1-Δ20 proteins were immobilized on the anti-His BLI biosensors and binding to Rpo41 was assessed by measuring the time-dependent increase in the light interference signal (Fig. 3A & S5). The amplitude of light interference corresponding to the amount of Rpo41-Mtf1 complex was recorded and plotted against increasing concentrations of Rpo41 (Fig. 3B & S5). The resulting binding curves indicated robust binding of Mtf1-WT to Rpo41 and relatively little binding of Mtf1-Δ20 to Rpo41.

**Fig. 3:**
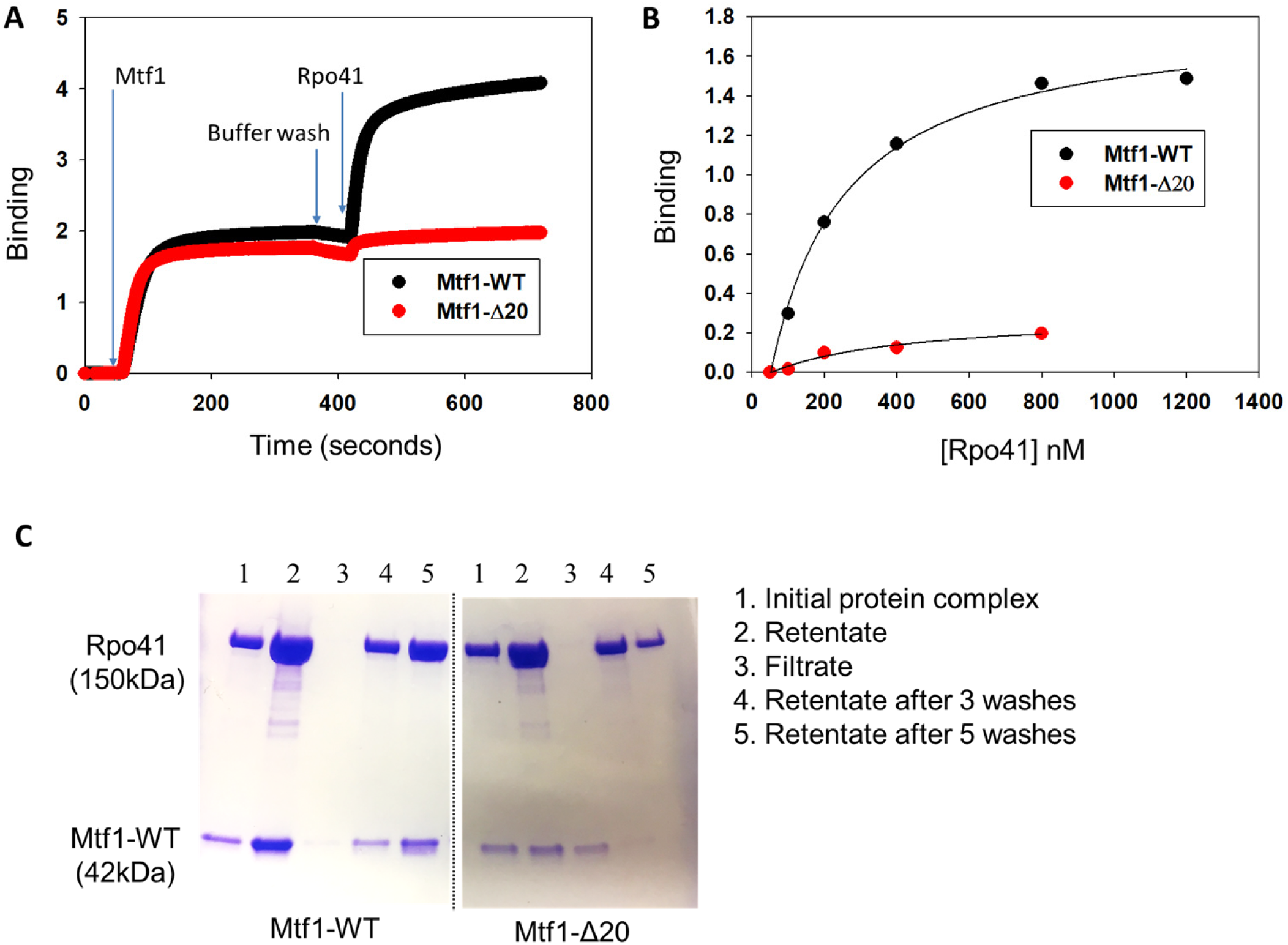
The C-tail of Mtf1 mediates complex formation with Rpo41. (A) Representative binding plots showing complex formation between Mtf1 and Rpo41 using Biolayer interferometry assays. The first 60 s represents the baseline. In the next 300s, the biosensor HIS1K was treated with 0.4 µM of His-tagged Mtf1-WT (black line) or Mtf1-Δ20 (red line) protein followed by washing with buffer for 60 seconds. The probes were then dipped in Rpo41 (0.5 µM) for 300 seconds followed by washing for 60 seconds. (B) The degree of binding (y-axis) was calculated from the difference in the light interference values before and after adding Rpo41 in (A). (C) An equimolar complex of Rpo41 and Mtf1-WT or Mtf1-Δ20 at a final concentration 2 µM (lanes 1) was filtered through a 100 kDa MW cut-off Microcon centrifugal filter unit. Lanes 2 are the rententates and lanes 3 are the filtrates. The retentate was washed with 500 µl of buffer 3 times (lanes 4). The retentate was washed two more times (lanes 5). Samples for initial protein complex, retentate, filtrate and retentate samples after washing were run on a 4-20% SDS-PAGE gel.

Ultrafiltration is an alternative solution-based method that has been used previously to assay complex formation between Rpo41 and Mtf1 (19). Application of this method also showed efficient complex formation between Rpo41 and Mtf1-WT as evident from retention of both proteins in the retentate (even after several washes) and almost no protein in the filtrate (Lanes 2 & 3 in Mtf1-WT in Fig. 3C). A stable complex was not observed with Mtf1-Δ20 (Lanes 2 & 3 in Mtf1-Δ20 in Fig. 3C). Rpo41 was present in the retentate and most of the Mtf1-Δ20 was in the filtrate.

The above methods of analysis of protein-protein interactions provide consistent results indicating that the C-tail of Mtf1 is involved in complex formation with the RNAP subunit.

### The C-tail of TFB2M promotes a stable initiation complex with POLRMT

We tried measuring complex formation between TFB2M and POLRMT in the absence of DNA using the BLI and the ultrafiltration methods, but these methods failed to provide consistent results, because TFB2M binds non-specifically to the biosensor probe and the filters. Hence, we used transcription assays and followed run-off products to monitor complex formation between TFB2M and POLRMT in the initiation complex.

We used TFB2M-Δ3 and TFB2M-Δ7 in the runoff assays, because TFB2M-Δ13 does not support transcription (3). Runoff synthesis was measured using *in vitro* reconstituted complex of POLRMT, TFB2M, TFAM, and the LSP promoter (Fig. 4A). Transcription initiation from the LSP DNA produces short abortive RNAs and two run-off products, 17- and 18-mer in length (Fig. 4B & S6). The two runoff products result from the two reported start-sites on LSP (underlined in the LSP sequence in Fig. 4A) (3,7).

**Fig. 4:**
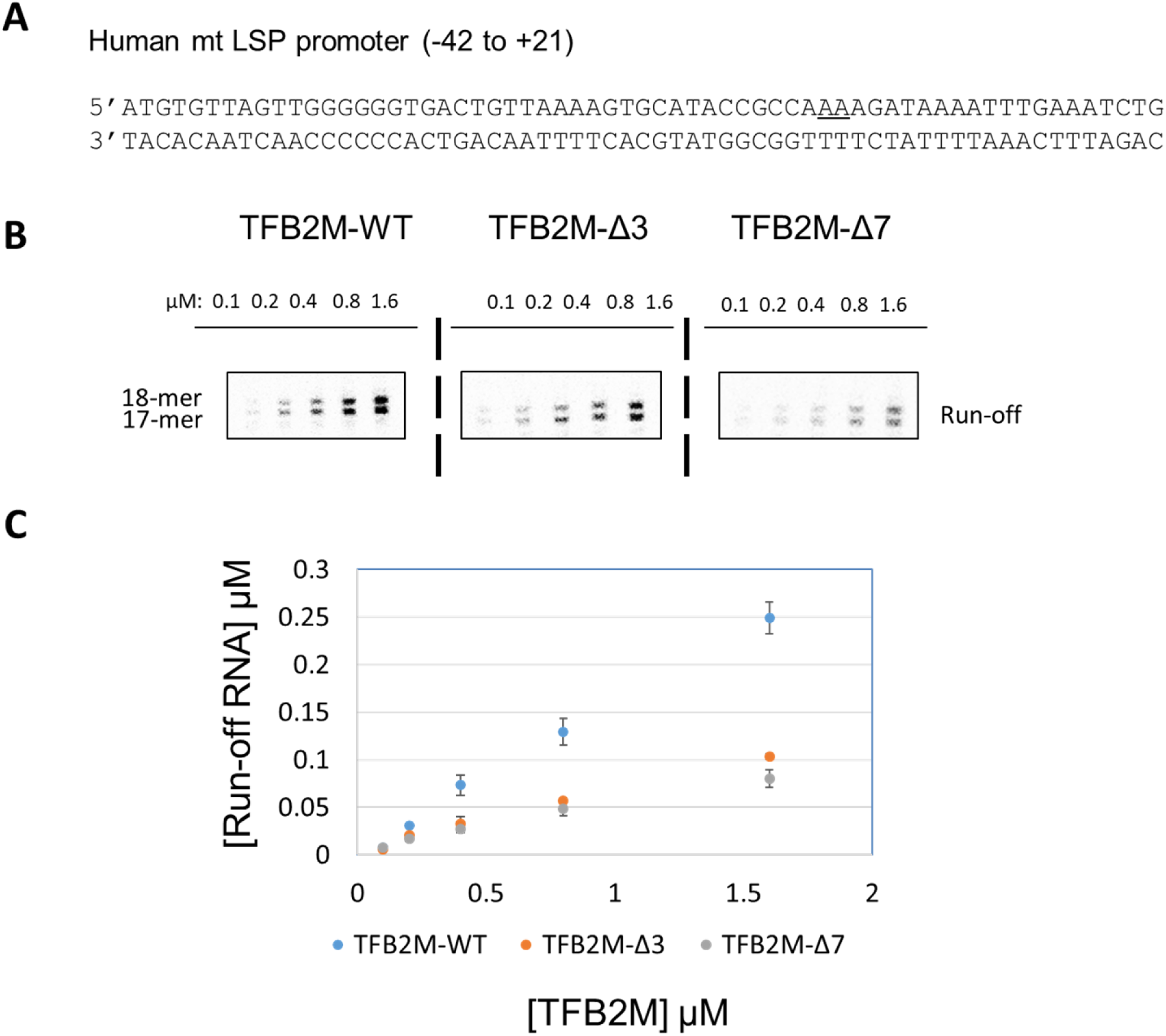
The C-tail deletion in TFB2M affects initiation complex formation. (A) The promoter fragment LSP used in this study is shown. The two start sites are underlined. (B) Run-off transcription profiles of TFB2M-WT and C-tail deletion mutants TFB2M-Δ3 and TFB2M-Δ7 on the LSP promoter are shown (The full gel profile is shown in Fig. S6). Reactions were carried out with 0.6 μM of POLRMT, TFAM, and promoter duplex and increasing concentration of TFB2M-WT or C-tail mutant and 250 μM ATP, UTP, GTP and γ[^32^P]ATP for 15 min at 25°C in transcription buffer. (C) Plot showing quantitation of grouped run-off products for each reaction. The error bars represent errors calculated from two independent experiments.

To measure the stability of the initiation complexes containing TFB2M-WT and C-tail deletion mutants, the transcription reactions were carried out at increasing TFB2M concentration (Fig. 4B-C & S6). TFB2M-WT reactions showed a steeper increase in runoff RNA products with increasing TFB2M concentration in comparison to TFB2M-Δ3 and TFB2M-Δ7. Close to 4-fold greater amounts of mutant proteins were required to observe the same amount of runoff products as with TFB2M-WT. These results indicate that the initiation complexes with the C-tail deletion mutants are weaker in comparison to TFB2M-WT. The results are consistent with the model that TFB2M C-tail is involved in complex formation with the RNAP subunit.

Thus, the results with Mtf1 and TFB2M suggest a consistent mechanism of activation of the DNA binding activity of the mitochondrial transcription factors.

## Discussion

Mitochondrial transcription factors, Mtf1 and TFB2M, of the yeast (*S. cerevisiae*) and human, respectively, have a well-established role in promoter melting during transcription initiation (8,11,17,20). These homologous transcription factors bind to the non-template strand of the transcription bubble to drive the promoter melting step. In the present study, we show that the DNA binding activity of the transcription factors is autoinhibited by their C-terminal tail regions prior to their association with the RNAP subunit. Both Mtf1 and TFB2M contain a flexible C-terminal tail region of 16 to 20 aa that we recently showed plays important roles in transcription initiation (3). The present study shows that the C-tail has additional roles in regulating the DNA binding activity of the free factors.

Parallel studies of the yeast and human mitochondrial transcription factors provided an opportunity to compare and contrast the mechanism of DNA regulation by the C-tail region. We find that the C-tail region of both Mtf1 and TFB2M has a conserved role in autoinhibiting the DNA binding activity of the free factors. Quantitation of DNA binding activity, however, shows that the C-tail of Mtf1 drastically inhibits DNA binding whereas the C-tail of TFB2M only partially inhibits DNA binding. Moreover, it appears that the DNA binding activity of the free factors is non-specific. The activated factors bind to both single-stranded and double-stranded DNA. Based on our results, we propose that Mtf1 is unlikely to associate with non-specific DNA regions prior to initiation. On the other hand, unless there are alternative means of inhibiting the DNA binding activity, the free TFB2M will associate with DNA prior to initiation.

Based on the crystal structure of TFB2M, we propose that the flexible C-tail can exist in two conformations, an autoinhibited state and a free-state (Fig. 5B). The crystal structure of free TFB2M shows two conformations of the C-tail in the two molecules of the asymmetric unit (Fig. 5A, cyan and white). In chain A, the C-tail (in red) is in the autoinhibited state making intramolecular interactions with the DNA binding groove and masking the DNA binding site. In chain B, the partially resolved C-tail (white) projects away from the DNA binding groove and more in line with the C-tail in the initiation complex (in green).

**Fig. 5:**
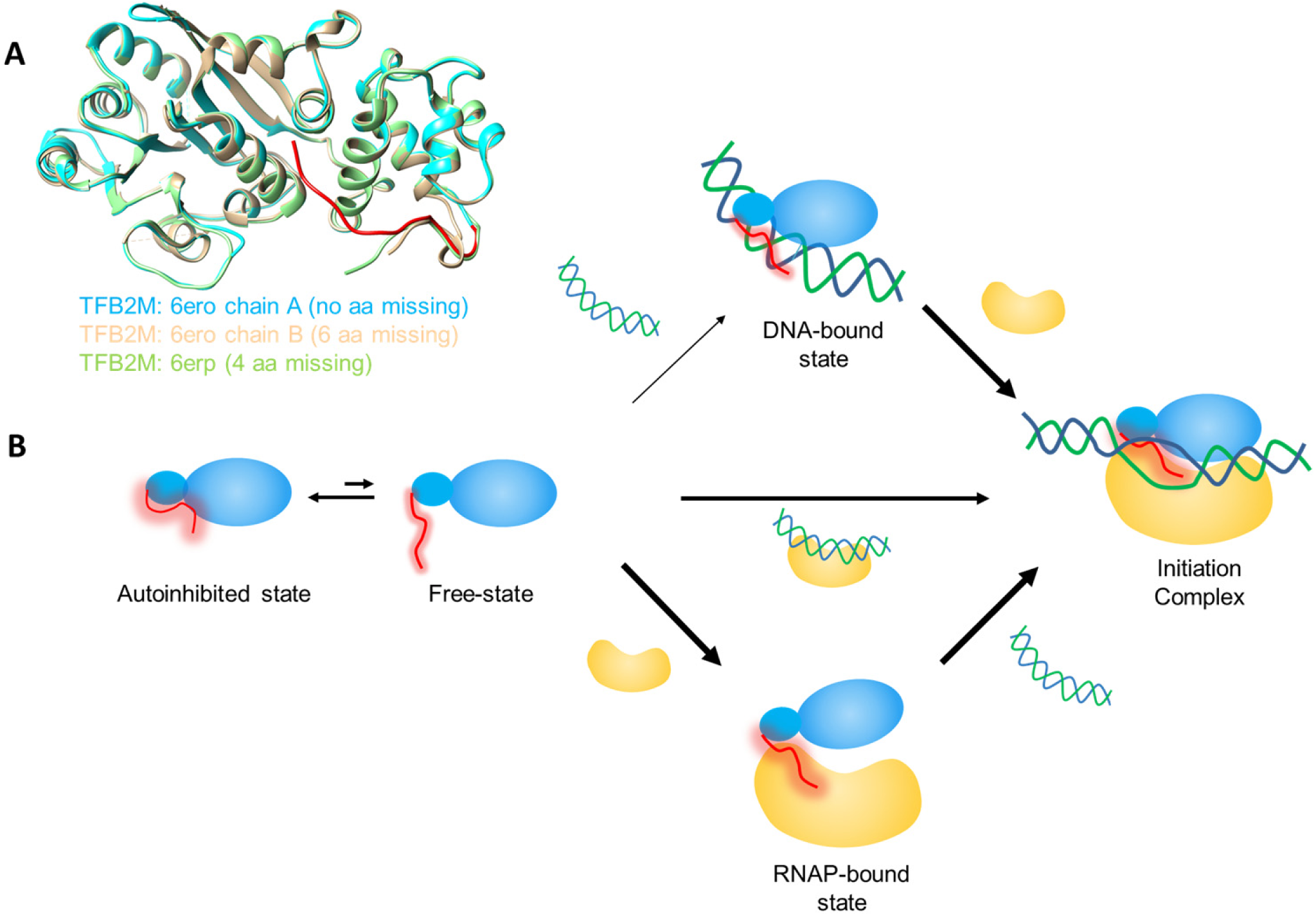
Model to explain the autoregulatory role of the C-tail of TFB2M and Mtf1 in the assembly of the initiation complex. (A) Crystal structures of TFB2M showing different states of the C-tail in free and DNA-bound states. Chain A of TFB2M in free state is shown in cyan and the C-tail in red, Chain B of TFB2M in free state is shown in white, and TFB2M in the initiation complex is shown in green. (B) The flexible C-tail (in red) of the mitochondrial transcription factors Mtf1 and TFB2M is in equilibrium between an autoinhibited state and free-state. For Mtf1, the equilibrium is towards the autoinhibited state, whereas for the TFB2M, both states exist. Therefore, Mtf1 binds to the RNAP or RNAP-DNA complex to generate the initiation complex. On the other hand, TFB2M can take all three pathways to form the initiation complex. Under cellular conditions, the DNA binding activity of TFB2M might be regulated by other mechanisms.

We propose that the equilibrium constant of the free and autoinhibited states is different in Mtf1 and TFB2M. The Mtf1 C-tail is mostly in the autoinhibited state whereas a significant portion of the TFB2M C-tail is in the free-state. It is not known if TFB2M binds DNA under cellular conditions, because it might be regulated by additional mechanisms. For example, it is known that TFB2M is post-translationally modified and that some of the modifications are in the C-tail region (13). Thus, the C-tail autoinhibition mechanism can be potentially reinforced by C-tail modifications. However, further studies are needed to understand these alternative mechanisms of TFB2M regulation.

Our studies also provide insights into how the transcription factors are activated for promoter-specific transcription initiation. It turns out that the C-tail region is essential for forming a stable complex with the RNAP. Thus, it appears that the RNAP subunit can activate the factors by engaging the C-tail with itself and thus releasing it from the autoinhibited state. The structure of TFB2M shows that the C-tail relocates from the DNA binding groove and gets engaged with the thumb domain and the intercalating hairpin of the RNAP in the initiation complex (Fig. S4) (8). The C-tail engagement with the RNAP would expose the DNA binding groove, which can bind to the non-template strand and stabilize the transcription bubble in the initiation complex.

Autoregulatory mechanisms are widely found in biological processes including DNA transcription. Bacterial and eukaryotic transcription factors such as sigma-70 and Ets-1, have an in-built regulatory domain that modulates their DNA binding activity prior to transcription. The DNA binding activity of sigma-70 is autoinhibited by the sigma-1.1 domain, and similar to mitochondrial transcription factors, autoinhibition is relieved when sigma-70 forms a complex with the RNAP subunit (16). Autoinhibition by a flexible C-terminal tail is also found in various DNA binding proteins. For example, the DNA binding activity of gp2.5 protein of bacteriophage T7 and bacterial SSB protein is autoinhibited by their acidic C-tails (14,15), which as we propose for the mitochondrial transcription factors, competes for the basic DNA binding cleft with the DNA.

Thus, our studies have identified a previously unknown and conserved role of the C-tail in regulating the DNA binding activity of yeast and human mitochondrial transcription factors. Such autoregulatory mechanisms increase the specificity of the transcription reaction and prevent transcription from occurring at non-promoter sites.

## Experimental procedures

### Nucleic Acid Substrates

Oligodeoxynucleotides were custom-synthesized with 5’-end fluorescein modification and purified by HPLC (Integrated DNA Technologies, Coralville, IA, USA). DNA concentration was determined from absorbance at 260 nm and the corresponding molar extinction coefficients. Complementary single-stranded DNAs were mixed in 1:1 ratio, annealed at 95°C for 1 min, and cooled over an hour to room temperature to construct the duplex DNA molecules.

### Protein purification

The expression and purification of Mtf1, TFB2M and the respective C-tail deletion mutant proteins were carried out as reported previously(3,7,21). The yeast proteins were stored in 50% and the human proteins were stored in 10% glycerol at −80°C. The molar concentrations of the proteins were determined in guanidium-HCl buffer from absorbance measurements at 280 nm and the respective molar extinction coefficients.

### Fluorescence anisotropy experiments to determine equilibrium constant K_d_ of protein binding to DNA

Fluorescence anisotropy-based titration experiments were carried out on Fluoro-Max-4 spectrofluorometer (Jobin Yvon-Spex Instruments S.A., Inc.) at 25°C. Fluorescein-labeled DNA (5 nM) was titrated with Mtf1-WT, TFB2M or the C-tail deletion mutants. Reaction buffer A (50 mM Tris acetate, pH 7.5, 50 mM potassium glutamate, 10 mM magnesium acetate, 1 mM DTT and 0.05% Tween-20) was used in studies of Mtf1, and Reaction buffer B (50 mM Tris-acetate pH 7.5, 100 mM sodium glutamate, 10 mM magnesium acetate, DTT 10 mM, 0.01% Tween-20) was used in studies of the TFB2M and. Anisotropy values (*r*obs) were recorded with excitation at 490 nm (4-nm bandwidth) and emission at 514 nm (8-nm bandwidth). The *r*obs was plotted against total protein concentration [P] and fit to Equation 1 to obtain the equilibrium dissociation constant (*K*_d_). Here rmax is the fluorescence anisotropy amplitude and rf is the initial fluorescence anisotropy of the DNA before protein addition.

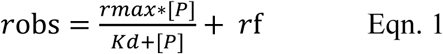

### Biolayer interferometry

The BLITZ binding assays were performed with Fortebio BLITZ system with a Dip and Read Penta-HIS (HIS1K) biosensor (ForteBio, CA, USA) using the tube method to measure complex formation between Rpo41+DNA and Mtf1. The probes were equilibrated in water overnight and then in Buffer A for 30 min prior to use. The biosensors were washed in Buffer A for 60 s to obtain the baseline. Mtf1 (400 nM) was immobilized on HIS1K biosensors for 300 s, washed in buffer A for 60 s, and then dipped in increasing concentration of Rpo41+DNA solution (50-800 nM) and finally washed again in Buffer A for 60 s. The wavelength shift was recorded in real-time with the ForteBio software.

### Ultracentrifugation assays

An equimolar complex of Rpo41 and Mtf1 (2 µM each) was mixed in Reaction buffer C (50 mM Tris acetate, pH 7.5, 100 mM potassium glutamate, 10 mM magnesium acetate, 5 mM fresh DTT, 0.01% protein-grade Tween 20, 5% glycerol) in a final volume of 500 µL. The mixture was incubated at 25 °C for 15 min (initial protein complex) before filtering through a 100 kDa MW cut-off Microcon centrifugal filter unit until the volume of the first retentate was about 50 µl (1/10 of initial mixture). The retentate diluted to 500 µl with Buffer C and filtered again. This washing step was repeated and a sample was taken after 3 and 5 washes. Samples consisting of initial protein complex, first retentate, filtrate and 3 and 5 retentate samples were collected and run on a 4-20% SDS-PAGE gel.

### Transcription reactions

Transcription reactions were carried out at 25°C using 1 µM POLRMT, 1 µM TFAM, 1 µM TFB2M-WT or the deletion mutants, and 1 µM of promoter DNA in transcription buffer (50 mM Tris-acetate pH 7.5, 100 mM sodium glutamate, 10 mM magnesium acetate, DTT 10 mM, 0.01% Tween-20). For runoff RNA synthesis, we used 250 μM ATP, UTP, GTP spiked with [γ-^32^P]ATP. Reactions were stopped after 15 min using 400 mM EDTA and formamide dye (98% formamide, 0.025% bromophenol blue, 10 mM EDTA). Samples were heated to 95°C for 2 min, chilled on ice, and the RNA products were resolved on 24% sequencing gel containing 4 M urea. The gel was exposed to a phosphor screen overnight and scanned on a Typhoon 9410 PhosphorImager instrument (Amersham Biosciences). The free ATP and RNA bands were quantified using ImageQuant, and molar amounts of RNA synthesized were calculated according to equation 2.

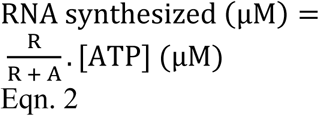

Where R and A are the band intensities of RNA products and free ATP, respectively, and [ATP] is the molar concentration of ATP added to the reaction.

## Data Availability

All the data are in the manuscript.

## Acknowledgements

We would like to thank the Patel Lab members for advice and suggestions on this work.

## Author contributions

U.B. and S.S.P. conceptualization; U.B., N.M., M.F., J.S. L.C.J. data curation; U.B. and S.S.P. formal analysis; U.B. and S.S.P. funding acquisition; U.B. and S.S.P. project administration; U.B. original draft, U.B. and S.S.P. writing and editing.

## FOOTNOTES

Funding for this work was provided by the American Heart Association (AHA) [16PRE30400001] and University and Louis Bevier Dissertation Completion Fellowship from Rutgers University to U.B. and the National Institute of General Medical Sciences (NIGMS) [GM118086 MIRA] to S.S.P.

## Figure Legends

**Fig. 1: The C-tail of Mtf1 drastically autoinhibits the DNA binding activity of Mtf1.** (A) Structure of the yeast mitochondrial transcription factor Mtf1 in grey (PDB ID:1i4w). The missing 16 aa of the C-tail of Mtf1 in the crystal structure are shown in red dotted line and also marked in red in the amino acid sequence of the C-tail of Mtf1. (B) DNA sequences of the substrates used for the Mtf1 DNA binding studies. (C) Cartoon showing the basic scheme of the fluorescence anisotropy assays to monitor protein-DNA binding. (D) Representative binding curves showing the fluorescence anisotropy changes resulting from titration of the *15S* NT DNA with Mtf1. *15S* NT DNA (5 nM) was titrated with Mtf1-WT (black circles), Mtf1-Δ12 (grey circles) and Mtf1-Δ20 (red circles) in Buffer A (Experimental procedures). (E) *15S* NT (5 nM) was titrated with Mtf1-WT (black circles), Mtf1-Δ12 (grey circles) and Mtf1-Δ20 (red circles) in Buffer A without potassium glutamate. The solid lines represent fit to the hyperbolic equation 1 with *K*_d_ values as follows: Mtf1-WT = 447±60 nM (amplitude: 0.059); Mtf1-Δ12 = 426±33 nM (amplitude: 0.156); Mtf1-Δ20 = 51±2.8 nM (amplitude: 0.24). The errors represent standard error of the fit. (F) The average DNA *K*_d_ values of Mtf1-Δ20 are shown for the DNA substrates in (B). The blue dots are the individual values for set 1 and set 2 titration data which are shown in Figure S1.

**Fig. 2: The C-tail of TFB2M mildly autoinhibits the DNA binding activity of TFB2M.** (A) The aligned structures of free human TFB2M (PDB: 6ero in cyan), TFB2M in the initiation complex (PDB: 6erp in green), and the yeast Mtf1 (PDB: 1I4W in white) are shown. The relative positions of the C-tail in all three structures are shown. The amino acid sequence of the C-tail of TFB2M is shown below in red. (B) DNA substrates used for the TFB2M DNA binding studies. (C) Representative binding curves show the fluorescence anisotropy change resulting from titration of LSP NT (5 nM) with TFB2M-WT (black circles), TFB2M-Δ3 (pink circles), and TFB2M-Δ13 (red circles). The data were fit to the hyperbolic equation 1 to obtain the following *K*_d_ values: TFB2M-WT = 169±18 nM (amplitude 0.17); TFB2M-Δ3 = 92±3.3 nM (amplitude 0.19); TFB2M-Δ13: 46±5.2 nM (amplitude 0.23). (D) The red and grey bars show the DNA *K*_d_ values of TFB2M-WT and TFB2M-Δ13, respectively, for the various DNA substrates shown in (B). The blue dots represent individual values for set 1 and set 2 titration data that are shown in Figure S3.

**Fig. 3: The C-tail of Mtf1 mediates complex formation with Rpo41.** (A) Representative binding plots showing complex formation between Mtf1 and Rpo41 using Biolayer interferometry assays. The first 60 s represents the baseline. In the next 300s, the biosensor HIS1K was treated with 0.4 µM of His-tagged Mtf1-WT (black line) or Mtf1-Δ20 (red line) protein followed by washing with buffer for 60 seconds. The probes were then dipped in Rpo41 (0.5 µM) for 300 seconds followed by washing for 60 seconds. (B) The degree of binding (y-axis) was calculated from the difference in the light interference values before and after adding Rpo41 in (A). (C) An equimolar complex of Rpo41 and Mtf1-WT or Mtf1-Δ20 at a final concentration 2 µM (lanes 1) was filtered through a 100 kDa MW cut-off Microcon centrifugal filter unit. Lanes 2 are the rententates and lanes 3 are the filtrates. The retentate was washed with 500 µl of buffer 3 times (lanes 4). The retentate was washed two more times (lanes 5). Samples for initial protein complex, retentate, filtrate and retentate samples after washing were run on a 4-20% SDS-PAGE gel.

**Fig. 4: The C-tail deletion in TFB2M affects initiation complex formation.** (A) The promoter fragment LSP used in this study is shown. The two start sites are underlined. (B) Run-off transcription profiles of TFB2M-WT and C-tail deletion mutants TFB2M-Δ3 and TFB2M-Δ7 on the LSP promoter are shown (The full gel profile is shown in Fig. S6). Reactions were carried out with 0.6 μM of POLRMT, TFAM, and promoter duplex and increasing concentration of TFB2M-WT or C-tail mutant and 250 μM ATP, UTP, GTP and γ[^32^P]ATP for 15 min at 25°C in transcription buffer. (C) Plot showing quantitation of grouped run-off products for each reaction. The error bars represent errors calculated from two independent experiments.

**Fig. 5: Model to explain the autoregulatory role of the C-tail of TFB2M and Mtf1 in the assembly of the initiation complex.** (A) Crystal structures of TFB2M showing different states of the C-tail in free and DNA-bound states. Chain A of TFB2M in free state is shown in cyan and the C-tail in red, Chain B of TFB2M in free state is shown in white, and TFB2M in the initiation complex is shown in green. (B) The flexible C-tail (in red) of the mitochondrial transcription factors Mtf1 and TFB2M is in equilibrium between an autoinhibited state and free-state. For Mtf1, the equilibrium is towards the autoinhibited state, whereas for the TFB2M, both states exist. Therefore, Mtf1 binds to the RNAP or RNAP-DNA complex to generate the initiation complex. On the other hand, TFB2M can take all three pathways to form the initiation complex. Under cellular conditions, the DNA binding activity of TFB2M might be regulated by other mechanisms.

